# Single cell transcriptomics reveals molecular subtype and functional heterogeneity in models of breast cancer

**DOI:** 10.1101/282079

**Authors:** Daniel L. Roden, Laura A. Baker, Benjamin Elsworth, Chia-Ling Chan, Kate Harvey, Niantao Deng, Sunny Z. Wu, Aurelie Cazet, Radhika Nair, Alexander Swarbrick

## Abstract

Breast cancer has long been classified into a number of molecular subtypes that predict prognosis and therefore influence clinical treatment decisions. Cellular heterogeneity is also evident in breast cancers and plays a key role in the development, evolution and metastatic progression of many cancers. How clinical heterogeneity relates to cellular heterogeneity is poorly understood, so we approached this question using single cell gene expression analysis of well established *in vitro* and *in vivo* models of disease.

To explore the cellular heterogeneity in breast cancer we first examined a panel of genes that define the PAM50 classifier of molecular subtype. Five breast cancer cell line models (MCF7, BT474, SKBR3, MDA-MB-231, and MDA-MB-468) were selected as representatives of the intrinsic molecular subtypes (luminal A and B, basal-like, and Her2-enriched). Single cell multiplex RT-PCR was used to isolate and quantify the gene expression of single cells from each of these models, and the PAM50 classifier applied. Using this approach, we identified heterogeneity of intrinsic subtypes at single-cell level, indicating that cells with different subtypes exist within a cell line. Using the Chromium 10X system, this study was extended into thousands of cells from the MCF7 cell-line and an ER+ patient derived xenograft (PDX) model and again identified significant intra-tumour heterogeneity of molecular subtype.

Estrogen Receptor (ER) is an important driver and therapeutic target in many breast cancers. It is heterogeneously expressed in a proportion of clinical cases but the significance of this to ER activity is unknown. Significant heterogeneity in the transcriptional activation of ER regulated genes was observed within tumours. This differential activation of the ER cistrome aligned with expression of two known transcriptional co-regulatory factors of ER (FOXA1 and PGR).

To examine the degree of heterogeneity for other important phenotypic traits, we used an unsupervised clustering approach to identify cellular sub-populations with diverse cancer associated transcriptional properties, such as: proliferation; hypoxia; and treatment resistance. In particular, we show that we can identify two distinct sub-populations of cells that may have *denovo* resistance to endocrine therapies in a treatment naïve PDX model of ER+ breast cancer. One of these consists of cells with a non-proliferative transcriptional phenotype that is enriched for transcriptional properties of ERBB2 tumours. The other is heavily enriched for components of the primary cilia. Gene regulatory networks were used to identify transcription factor regulons that are active in each cell, leading us to identify potential transcriptional drivers (such as E2F7, MYB and RFX3) of the cilia associated endocrine resistant cells. This rare subpopulation of cells also has a highly heterogenous mix of intrinsic subtypes highlighting a potential role of intra-tumour subtype heterogeneity in endocrine resistance and metastatic potential.

Overall, These results suggest a high degree of cellular heterogeneity within breast cancer models, even cell lines, that can be functionally dissected into sub-populations of cells with transcriptional phenotypes of potential clinical relevance.

## Introduction

Breast cancer molecular subtype is used to guide patient treatment and is crucial in determining a patients’ predicted responsiveness to targeted therapies (Perou et al., 2000; Sorlie et al., 2001). The determination of intrinsic molecular subtype from bulk patient tumour RNA samples has become an important clinically useful diagnostic tool to predict a patient s risk of relapse and response to chemotherapy (Ayers et al., 2004; Perou et al., 2000; Sorlie et al., 2001). However, patients do not present with homogenous tumours, and often lack uniform expression of crucial therapeutic targets including Estrogen receptor (ER) and human epidermal growth factor receptor (Her2) (Greer et al., 2013; Ng et al., 2015; Perou et al., 2000; Seol et al., 2012; Shimada et al., 1985; Sorlie et al., 2001) and reviewed in (Rivenbark, O’Connor, & Coleman, 2013). This heterogeneity impacts clinical assessment of the tumour and the application of targeted treatment strategies and may facilitate recurrence (Parker et al., 2009; Potts et al., 2012; Sorlie et al., 2001). To better diagnose and treat patients, we may need to accurately assess intra-tumoural heterogeneity by studying intrinsic molecular subtype at the cellular level.

The power of single-cell analysis is revolutionising biological research. Researchers have compared gene expression signatures, focal copy number variation and whole-chromosome aneuploidy, inferring evolutionary relationships both within primary tumours and with distant metastases (Navin et al., 2011). Single-cell technologies are enabling detailed analysis of samples with heterogeneous cell cycle stages, mutation profile and single-nucleotide polymorphisms (SNPs) (Wills et al., 2013). The clinical implications of these findings are likewise emerging, with remarkable insights being gained from single-cell studies on the impact of intra-tumoural gene expression and cancer subtype heterogeneity in a number of cancer types (Patel et al., 2014; Tirosh et al., 2016). Recent technological advances have massively increased the number of cells that can be profiled in a single experiment, making the generation of transcriptional maps of 1000’ s of cells now possible (Macosko et al., 2015; Zheng et al., 2017). Although technological and computational challenges are still to be solved, these methods and orthogonal approaches (Stoeckius et al., 2017) are expected to revolutionise our understanding of the cellular ecosystems present in complex diseases like cancer.

Recent single-cell studies in breast cancer have shown that transcriptional heterogeneity can be characterised using a number of different techniques and tumour cell types, such as: cell-lines; PDX models and primary clinical samples (Chung et al., 2017; Gao et al., 2017; Savage et al., 2017). Here, we use complementary single-cell technologies to explore the transcriptional properties that determine the complex intra-tumoural heterogeneity that is observed histologically in the malignant, neoplastic cells of breast cancers. To address this we selected several models of breast cancer to investigate the gene expression signatures of single-cells. These models include commonly studied breast cancer cell-lines and patient derived xenografts (PDXs). Breast cancer cell lines and PDXs have been extensively used to study and model cancer, enabling pre-clinical characterisation of drug response and evaluation of complex signalling pathways (Lacroix & Leclercq, 2004; Marangoni et al., 2007; Neve et al., 2006). PDX resources often reflect the degree of heterogeneity observed in the original patient tumour and recapitulate similar growth kinetics, metastatic potential and histopathological characteristics (Bruna et al., 2016; DeRose et al., 2011) and reviewed in (Tentler et al., 2012).

In this study we use these to model the gene expression signatures driving both intrinsic molecular subtype classifications and the heterogenous activation of transcriptional modules in cellular subpopulations of a tumour. Using single cell transcriptomics and the PAM50 intrinsic molecular subtype predictor, we demonstrate that tumours have intrinsic subtype heterogeneity at the single cell level, which is reflected in cell line and PDX models. We then show that, in ER+ disease, differential activation of the ER cistrome occurs in sub-populations of cells that correlate with coexpression of key co-regulatory factors of ER. In addition, unsupervised clustering is able to identify a sub-population of cells that show increased expression of genes associated with treatment resistance, highlighting cells that show putative, *de-novo* resistance to therapy in a treatment naïve setting.

## Results

### Breast cancer cell-lines show significant transcriptional heterogeneity at cellular resolution using a panel of breast cancer genes

To explore cellular heterogeneity in breast cancer we first used single cell microfluidics and multiplex RT-PCR to analyse expression of a selected panel of 96 genes. These included: the 50 genes comprising the PAM50 intrinsic breast cancer subtype predictor (Parker et al., 2009); genes that are markers of stem, mammary stem, luminal, basal and epithelial breast cancer cells as well as ER, Notch, Hedgehog and NF-kB signalling; and two house keeping genes (HKG) GAPDH and HPRT. We initially used five commonly studied breast cancer cell-lines to represent the intrinsic breast cancer molecular subtypes: MCF7 (Luminal A), BT474 (Luminal B), SKBR3 (Her2-E), MDA-MB-231 (Basal-like/Claudin-low) and MDA-MB-468 (Basal-like). A total of 359 single cells that successfully passed stringent QC criteria, for both cell viability and the presence of single cells, were identified for further analysis.

A subset of marker genes were selected to assess the ability of the method to characterise specific cell-lines and also gauge the heterogeneity across individual cells from the same models (Figure 1A and Supplementary fig 1). These genes were selected based on their importance to: ER signalling (ESR1, PGR, FOXA1); ERBB2 signalling (ERBB2); basal-like properties (KRT5); EMT (SNAI2); and proliferation (MKI67). The cell-lines clearly cluster based on expression of key lineage markers of breast cancer, with ESR1 and PGR restricted to the two ER+ cell-lines (BT and MCF7), ERBB2 expression in the SKBR3 cell-line (as well as BT), and the 468 and 231 ER-cell-lines showing expression of KRT5 and SNAI2, respectively (Figure 1A and Supplementary material). However, even this small subset of selected genes starts to highlight the presence of transcriptional heterogeneity of single-cells within a cell-line. In MCF7 cells, only a subset of cells have detectable PGR expression and, to a lesser extent, a subset of BT cells have no detectable expression of ESR1. We also see heterogenous expression of a breast cancer stem cell marker, CD44 (Al-Hajj, Wicha, Benito-Hernandez, Morrison, & Clarke, 2003), in subsets of cells from the MCF7 and BT cell-lines as well as robust expression in the two basal-like models (231 and 468) (Supplementary material).

**Figure 1.**
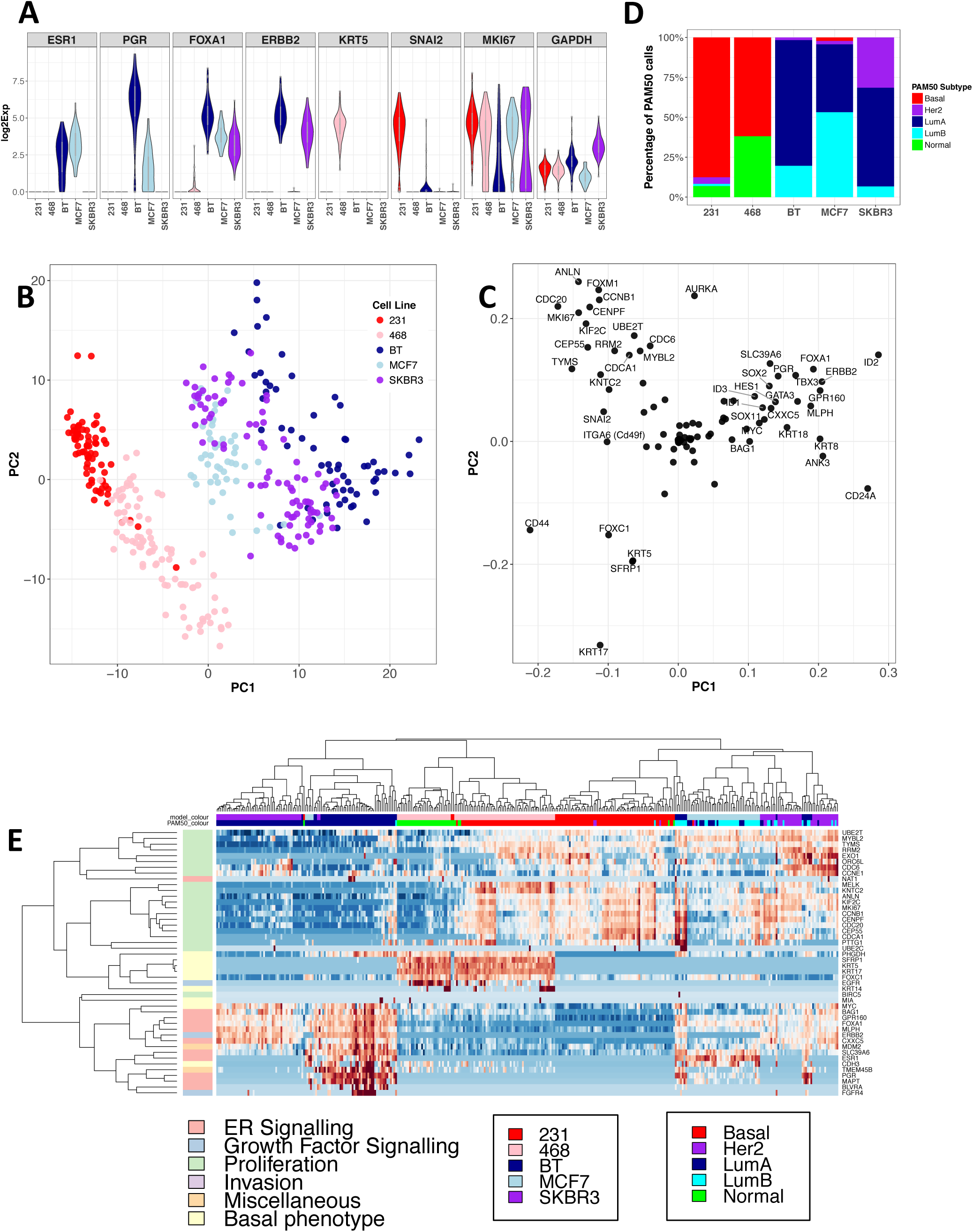
Breast cancer cell-lines show significant transcriptional heterogeneity at cellular resolution using a qPCR targetted panel of key breast cancer genes. **A)** Distribution of single-cell qPCR gene expression levels of selected breast cancer marker genes in five breast cancer cell lines. **B)** PCA plot of the first two principal components (PC1 and PC2) showing the clustering of the profiled cells from five breast cancer cell lines. **C)** PC gene loadings of the same cells shown in B. **D)** The percentage of PAM50 subtype calls for each cell across the five breast cancer cell lines. **E)** Heatmap showing hierarchical clustering of PAM50 gene expression levels from all cancer cells in the five cell-line models.

Extending this analysis to include all profiled genes, principle component analysis (PCA) shows that cell lines cluster into heterogeneous groups (Figure 1B) with a separation between the ER-models (231 and 468) and the others. The PCA loadings (Figure 1C) show that this separation is predominantly driven by stem, mesenchymal and basal-like genes (E.g. CD44, SNAI2, ITGA6); and luminal and ER signalling genes (E.g. CD24, KRT8, PGR, FOXA1). PCA analysis also highlights heterogeneity for proliferation, where the cells in the top-left region of the PCA plot (Figure 1B) are being driven by a set of proliferation genes (such as: MKI67, AURKA and KIF2C) (Figure 1C). These cells largely consist of the 231 and MCF7 and proliferative sub-populations of cells from the other 3 cell-lines.

### Individual breast cancer cell-lines show heterogenous distributions of intrinsic molecular subtype

To investigate how this observed transcriptional heterogeneity may impact on clinical markers of breast cancer we utilised the PAM50 method (Parker et al., 2009), which can determine breast cancer subtype and risk of recurrence. It was not known whether individual cells possess a discernible PAM50 phenotype, or whether it was dependent on a compendium of cellular states in bulk. Figure 1D clearly shows that there is considerable heterogeneity of PAM50 subtype calls for the cells within each of the cell-lines. This revealed some surprising findings, for example in the Skbr3 model (Her2-enriched in bulk), only a minority of cells (∼30%) were called as Her2-enriched, with the remainder luminal.

To better understand the transcriptional features driving this subtype heterogeneity we hierarchically clustered both the cells and the PAM50 genes (Figure 1E). This shows a substantial degree of intra-model subtype heterogeneity, with a major component driven by the proliferative capacity of the constituent cells. A more detailed analysis highlights six clusters of cells that have the following transcriptional characteristics: (i) proliferative with ER and growth factor signalling (far right) with predominantly HER2 and LumB subtype calls and a heterogenous mix of BT, MCF7 and SKBR3 cells; (ii) proliferative with low ER, growth factor or basal marker expression (these are mostly from the 231 model and have basal-like PAM50 calls); (iii) proliferative with strong expression of basal markers and low levels of ER and growth factor signaling (these are from the 468 model and are mostly classified as basal-like); (iv) non-proliferative with strong expression of basal markers and low ER and growth factor signaling (these are mostly normal-like cells from the 468 model); (v) non-proliferative with strong ER and ERBB2/growth factor signaling (which are mostly LumA cells from the BT-474 model); and finally (vi) non-proliferative with ERBB2 expression and some ER signaling components (such as FOXA1) but no ESR1 or PGR expression (these are from the SKBR3 model and are also classified as LumA).

### High-throughput analysis of breast cancer cells reveals cellular subpopulations of intrinsic molecular subtype and proliferative potential

Recent technological advances in the capture and transcriptional profiling of single-cells have made possible high-throughput, unbiased whole transcriptome profiling of 1000’ s of cells (Macosko et al., 2015; Zheng et al., 2017). We used the Chromium system from 10X Genomics (Zheng et al., 2017) to explore the transcriptional heterogeneity in cancer cells from two ER+ models of breast cancer: the MCF7 cell-line; and a treatment naïve PDX model of ER+ breast cancer (HCI003) (DeRose et al., 2011). These methods reduce both the potential sampling limitations, when trying to infer population level heterogeneity from small numbers of cells, and extend the transcriptional coverage to all mRNAs rather than a targeted gene panel approach.

A summarised gene count matrix was generated for each of the breast cancer models, resulting, after QC, in 1699 cells from MCF7 and 1927 cancer cells from the PDX. These datasets were then individually log normalized using Seurat (Satija, Farrell, Gennert, Schier, & Regev, 2015). We first repeated our analysis of PAM50 molecular subtype heterogeneity. Due to the lower diversity of datasets profiled with the high-throughput approach and that we analysed each dataset separately, it was decided that the subtype specific centering method of PAM50 classification was most appropriate (Zhao, Rødland, Tibshirani, & Plevritis, 2015). Figure 2A shows the distribution of PAM50 subtypes in each of these models. The results from the MCF7 cell-line show comparable distributions of subtype to that from the qPCR analysis, i.e., a majority of cells are classed as either Luminal A or B, with smaller numbers of cells classified as basal-like, Her2 and normal-like.

**Figure 2.**
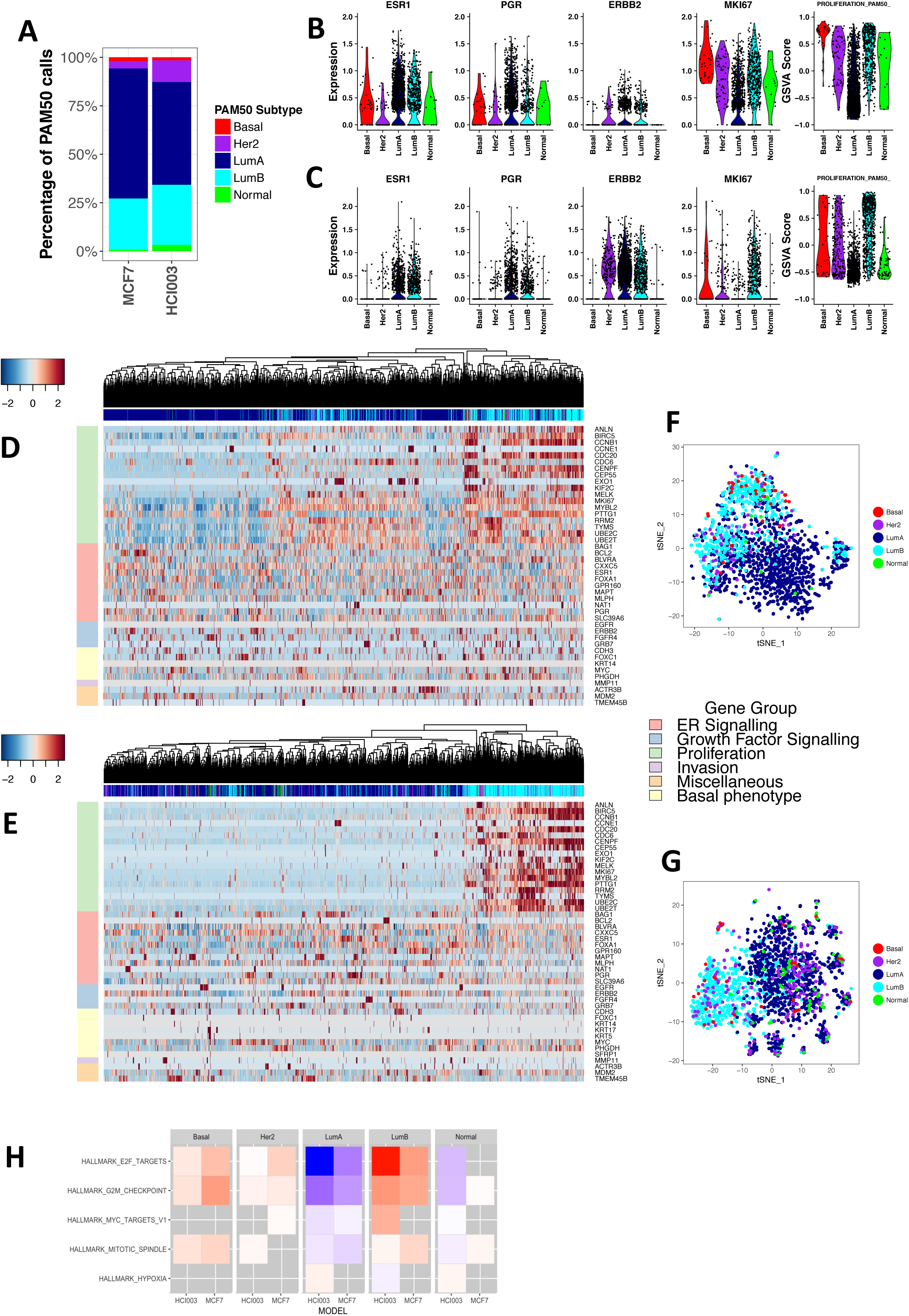
High-throughput 10X single-cell RNA-Seq of ER+ breast cancer cells reveals cellular subpopulations of intrinsic molecular subtype and functional heterogeneity. **A)** The percentage of PAM50 subtype calls for each cell across the three breast cancer models. **B-C)** Distribution of transcript expression levels for key breast cancer marker genes (ER, PR and ERBB2), proliferation marker (MKI67) and PAM50 proliferation gene signature score for individual cells stratified by their PAM50 molecular subtype classification, in MCF7 cells **(B)** and the PDX-HCI003 **(C)** models, respectively. **D and E)** Heatmaps showing hierarchical clustering of PAM50 gene expression levels from all cancer cells in the MCF7 **(D)** and PDX-HCI003 **(E)** models. The PAM50 subtype classification for each cell is shown at the top of the heatmaps and is the same as that in **(A). E and F)** t-SNE visualisations, generated from the gene expression level of the PAM50 genes, showing the clustering of all cancer cells in the MCF7 **(E)** and PDX-HCI003 **(F)** models. Each point represents a single-cell and are coloured according to their PAM50 subtype classification. **H)** Heatmap showing the five Hallmark gene signatures that are most enriched (|-log10(q-value)|) for the genes that positively mark each of the PAM50 subtypes in the two ER+ models of breast cancer.

To further validate the PAM50 classifications of the cells in these datasets a more detailed analysis of the expression of selected key marker genes (such as: ESR1, PGR, ERBB2 and MKI67) and functional signatures (such as proliferation, luminal, basal-like and HER2 signalling) was done to identify known association with the intrinsic subtype classifications of the cells. Specifically, we see higher expression of ESR1 and PGR (as predicted) in the cells classified as luminal A and B (figures 2B and 2C). Also, ESR1 and PGR expression appear higher in the Luminal A cells, compared to Luminal B, in both the cell-line and PDX. However, only PGR in the MCF7 cells has a significant difference (p=6.68e-12). ERBB2 is expressed in both the luminal and Her2 enriched cells, whereas the Her2 enriched cells have significantly reduced expression of both ESR1 and PGR. Increased proliferation is a known property of basal-like, luminal B and (to a lesser extent) Her2-enriched tumours. We can observe this same correlation at the single cell level, where the proliferation marker MKI67 and the PAM50 proliferation signature score are more highly expressed in the luminal B, basal-like and Her2 cells in both the ER+ models (figures 2B and 2C). Hierarchical clustering of the cells, using the expressed PAM50 genes, also highlights that subtype heterogeneity is strongly associated with the expression of proliferative genes. Specifically, in the malignant cells from the HCI003 PDX tumour, the Luminal B cells cluster together and appear to form a distinct group of proliferative cells (Figure 2E). t-SNE plots using the PAM50 genes also highlight these clear clusters of luminal B cells (figures 2E and 2G).

### Transcriptional properties of the PAM50 subtypes can be identified at cellular resolution

We next examined what other properties may differ between cells of varying intrinsic subtype, within a single tumour. For each model, genes that were positively and negatively enriched in each subtype (compared to the rest) were identified along with enriched functional gene signatures. As expected, the top-5 hallmark gene signatures in both datasets show a strong and significant positive enrichment for cell cycle and proliferation gene signatures (E2F_TARGETS, G2M_CHECKPOINT) in the luminal B, basal and (to a lesser extent) the Her2-enriched cells, with a corresponding negative enrichment in the luminal A cells (figure 2H).

In addition to these common transcriptional properties there are also a number of functional signatures that are restricted to cells in the individual models. This can be seen in the Hallmark analysis, where hypoxic gene expression is increased in the luminal A cells from the ER+ PDX but not the MCF7 cell-line, suggesting an interaction of intrinsic subtype with features of the tumour microenvironment (TME) *in vivo.* Extending the analysis to include all gene-sets in the MSigDB C2 CGP collection highlights a number of differentially enriched gene-sets that are of clinical relevance in ER+ breast cancers. In particular, both the Her2-enriched and basal-like cells from the ER+ PDX show positive enrichment for genes associated with endocrine therapy resistance. These also show decreased expression of genes that are down-regulated upon acquired endocrine resistance (MASSARWEH_ENDOCRINE_RESISTANCE_DN) and also the group 4 set of Creighton endocrine therapy resistance which were found to increase in models of endocrine therapy resistance (Creighton et al., 2008; Massarweh et al., 2008). This broadly agrees with their findings that tamoxifen resistance is associated with Her2 activation and highlights the potential role that intrinsic subtyping of individual tumour cells could play in identifying cells that may acquire or display *de-novo* resistance to therapy.

### Single-cell analysis provides a high-resolution map of cis- and co-regulatory action of hormone receptor signaling in ER+ breast cancer

Our observations in figure 2 suggest heterogenous expression of ER and its regulatory targets, such as PGR. To explore this further we extended our analysis of transcriptional heterogeneity to look in more depth at estrogen driven regulation in ER+ breast cancer cells. The MCF7 cell-line and other models of ER+ breast cancer have been extensively studied and have provided detailed understanding of the molecular drivers of estrogen regulated transcription and also the transcriptional targets of ER (Hurtado, Holmes, Ross-Innes, Schmidt, & Carroll, 2011). Using knowledge derived from gene expression and ChIP-Seq studies in bulk populations of MCF7 cells, we looked to develop a high-resolution cellular map of estrogen receptor driven transcriptional regulation.

We first hypothesised that the ER expression level within each cell should show positive correlations with: (i) the expression of the genes activated by estrogen; and (ii) those that are also ER ChIP-Seq targets (direct ER targets). This was indeed found to be the case in MCF7 cells, with moderate positive correlations of R=0.08 and R=0.17 observed for the estrogen induced and the direct ER target genes, respectively (Supplementary material). The cells were also split into those with any detected expression of ESR1 transcript and those without, with a significant enrichment (p=3.5e-6) for expression of ER target genes in those cells with detected ER transcripts (Supplementary material). The same analysis was carried out with the genes repressed by estrogen and ER but no negative correlation or significant depletion of gene expression in cells expressing ER was observed (Supplementary material). A similar and more significant result was seen in the cells from the ER+ PDX tumour, with a positive correlation of 0.19 and a significant difference (p<2.2e-16) between expression of ESR1 and the ER cis-regulatory targets (figure 3A).

**Figure 3.**
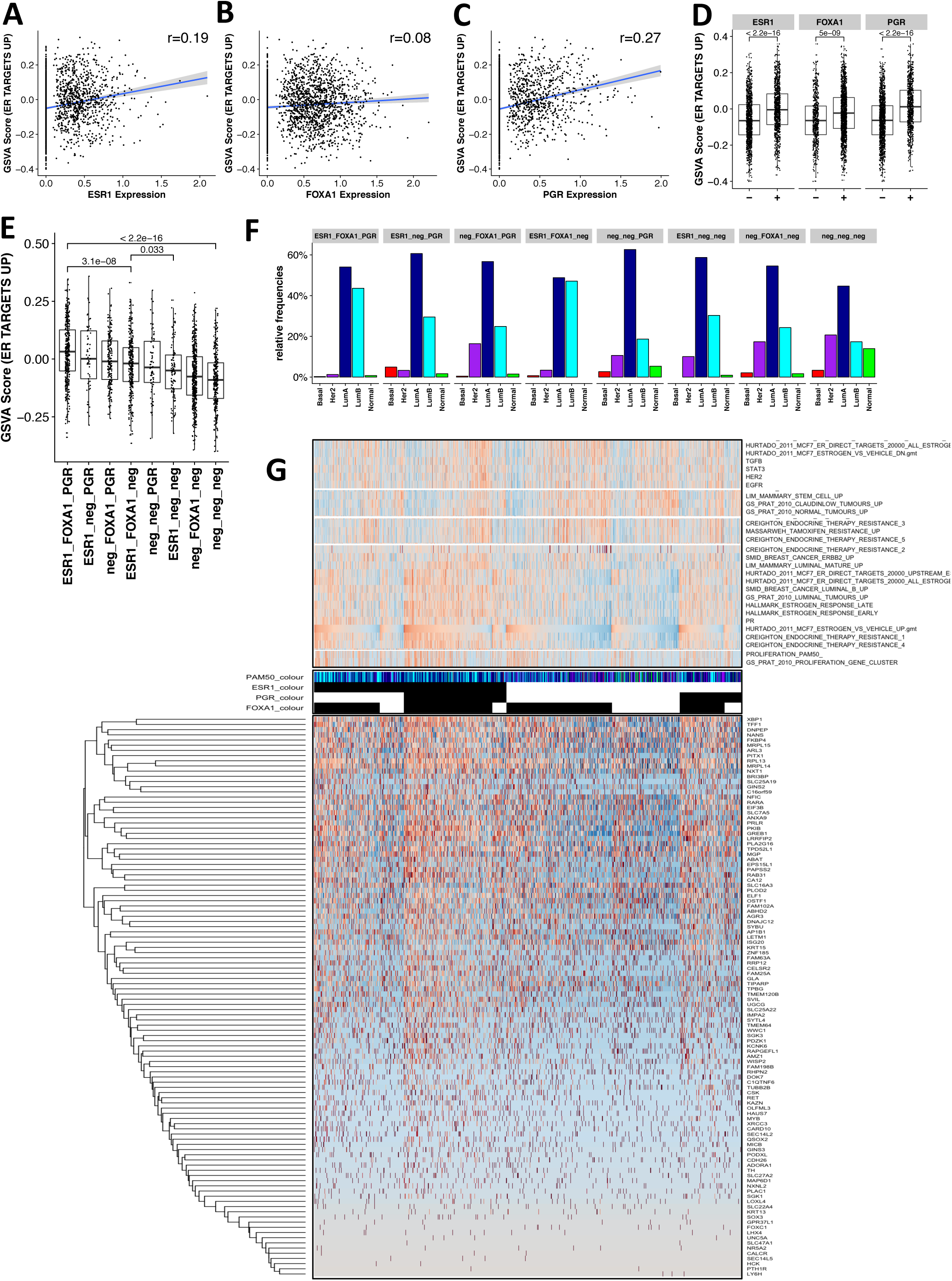
Single-cell analysis reveals heterogenous ER co-regulatory signalling and cis-regulatory action within ER+ breast cancer. **A-C)** Correlation between ESR1 **(A),** FOXA1 **(B),** PGR **(C)**expression and estrogen induced ER ChIP targets. **D)** Boxplots of signature scores of estrogen induced ER ChIP targets for each cell after stratification of cells into groups that have detectable expression (+) or not (-) of ESR1, FOXA1 or PGR. p-values calculated using Wilcox rank sum test. **E)** Boxplots of signature scores of estrogen induced ER ChIP targets after stratification of cells into groups defined by combinations of detectable expression of ESR1, FOXA1 and PGR. Boxplots ordered by median (descending). p-values calculated using Wilcox rank sum test. **F)** Relative frequency of PAM50 subtypes in each of the eight cell groupings. **G)** Heatmap of row-scaled GSVA signature scores from selected gene-signatures (bottom) and row clustered heatmap of estrogen induced ER ChIP target genes (bottom). Cells are grouped depending on whether there is detectable expression of ESR1, FOXA1 or PGR. Within each of the 8 gene expression based groupings the cells are ranked in descending order according to their estrogen induced ER ChIP target GSVA score.

We next looked at the effect of two known co-regulatory factors of ER modulation (FOXA1 (Hurtado et al., 2011) and PGR (Mohammed et al., 2015)) on the transcriptional activation of direct ER target genes. These co-regulators are also of interest as they are defining genes of luminal breast cancers and part of the PAM50 classifier. As with ER, we looked at the association of FOXA1 and PR expression with activation of all estrogen activated genes and the direct ER ChIP targets. In general, the transcriptional signal and heterogeneity was clearer in the PDX tumour cells and as such we concentrate our discussion on the results from these cells (results from the MCF7 cells are shown in supplementary material). Expression of the pioneer factor, FOXA1, was also found to have a significant association with estrogen induced (R=0.11 and p<2.2e-16) and direct ER target gene expression (R=0.08 and p=5.0e-09, figures 3B and 3D). This result is in contrast to MCF7 where the association with FOXA1 expression is insignificant. Similarly, the expression of PR showed a significant and positive correlation with the ER regulated gene signatures (figures 3C and 3D). This agreed with expectation as PR is one of the genes significantly up-regulated by estrogen stimulation in MCF7 cells. Interestingly, correlation of PR expression with direct ER target gene expression (R=0.27, figure 3C) was higher than ER itself (R=0.19, figure 3A).

We next reasoned that if the cells were grouped on whether they expressed each (or a combination) of these three transcription factors we should see differential activation of the ER driven genes depending on their patterns of co-expression. This was indeed the case, with a significant enrichment (p<2.2e-16) for ER target genes in the cells expressing all three genes compared to those cells with no detectable expression of any (figure 3E). Perhaps surprisingly, ER expression alone, or the co-expression of only ER and FOXA1, wasn’t sufficient to “maximally activate” the ER target genes. However, significantly increased activation was observed when all three were expressed in the same cell, compared to those cells only expressing ER and FOXA1 (p=3.1e-08) (figure 3E). This suggests that the strongest activation of ER target genes occurs when all 3 factors are present in the one cell. This pattern of heterogenous ER-driven gene expression, within each of the eight cell populations identified in figure 3E, can be clearly seen in figure 3G. In addition, a negative correlation is identified between cells that have activation of the E2 induced genes (HURTAD0_2011 ESTROGEN_vs_VEHICLE_UP) and those that have activation of genes that are repressed by E2 (HURTAD0_2011_ESTR0GEN_vs_VEHICLE_DN) (figure 3G, top).

An important and understudied aspect of ER+ breast cancer is the transcriptional features of cells that do not express ER (or any of its key co-regulatory factors such as PR and FOXA1). Our data allows us to provide a refined, high-resolution transcriptional map of these “receptor negative” populations of cancer cells in the context of heterogenous ER+ expression. We approached this by analysing the molecular subtypes assigned to the cells within each of the ER and co-factor related sub-groups (figure 3F) as well as the enrichment and correlation of ER target expression with selected gene signatures of functional and clinical interest (figure 3G, top). The cell subpopulations with the highest (ESR1_FOXA1_PGR) and lowest (neg_neg_neg) activation of ER targets also show a large difference in the level of subtype diversity, where the cells expressing all 3 factors are predominantly luminal cells (with >40% proliferative Luminal B cells) whilst the “all negative” cells have the lowest fraction of luminal A cells and similar proportions of Her2, LumB and normal-like classified cells (figure 3F). In addition, these cells have fewer proliferative cells and an increased enrichment for genes defining mammary stem cells and claudin-low tumours (figure 3G, top). The other subset of “receptor negative” cells, that express only FOXA1, also show a relatively high proportion of cells with the HER2-enriched subtype (figure 3F) and enrichment for transcriptional signatures found to be up-regulated in endocrine therapy resistant models of ER+ breast cancer (i.e., CREIGHTON_ENDOCRINE THERAPY groups 2, 3 and 5 and MASSARWEH_TAMOXIFEN_RESISTANT_UP) (Creighton et al., 2008; Massarweh et al., 2008)(figure 3H).

### Unsupervised clustering of single-cell RNA-Seq identifies sub-populations of functional heterogeneity and putative endocrine treatment resistant cells in a clinical model of ER+ breast cancer

We hypothesized that unsupervised methods should be able to identify functionally relevant subsets of cells and provide novel insights into the transcriptional properties and drivers of intratumour heterogeneity in breast cancer. Focusing on the PDX model, we identified 8 cellular subpopulations, “clusters”, across the ∼2000 profiled cancer cells (figure 4A). Expression of beta-actin (ACTB) was present and relatively consistent across all of the clusters (figure 4B), whereas expression of the breast cancer receptor marker genes (ESR1, PGR and ERBB2) was generally lower with a sparser, more heterogenous profile not clearly restricted to a specific cluster (figure 4B). We again looked for association between these unsupervised clusters and the molecular subtypes (Figures 4C and 4D). Luminal B cells are clearly enriched in cluster 2 (figure 4C and D), which is characterized by the more proliferative cancer cells that have significantly higher expression of MKI67 (figure 4B) and are marked by genes that are significantly enriched in proliferative and cell-cycle related gene-sets (figures 4E and 4F). In contrast, the luminal A cells are almost entirely absent in this proliferative cluster of cells.

**Figure 4.**
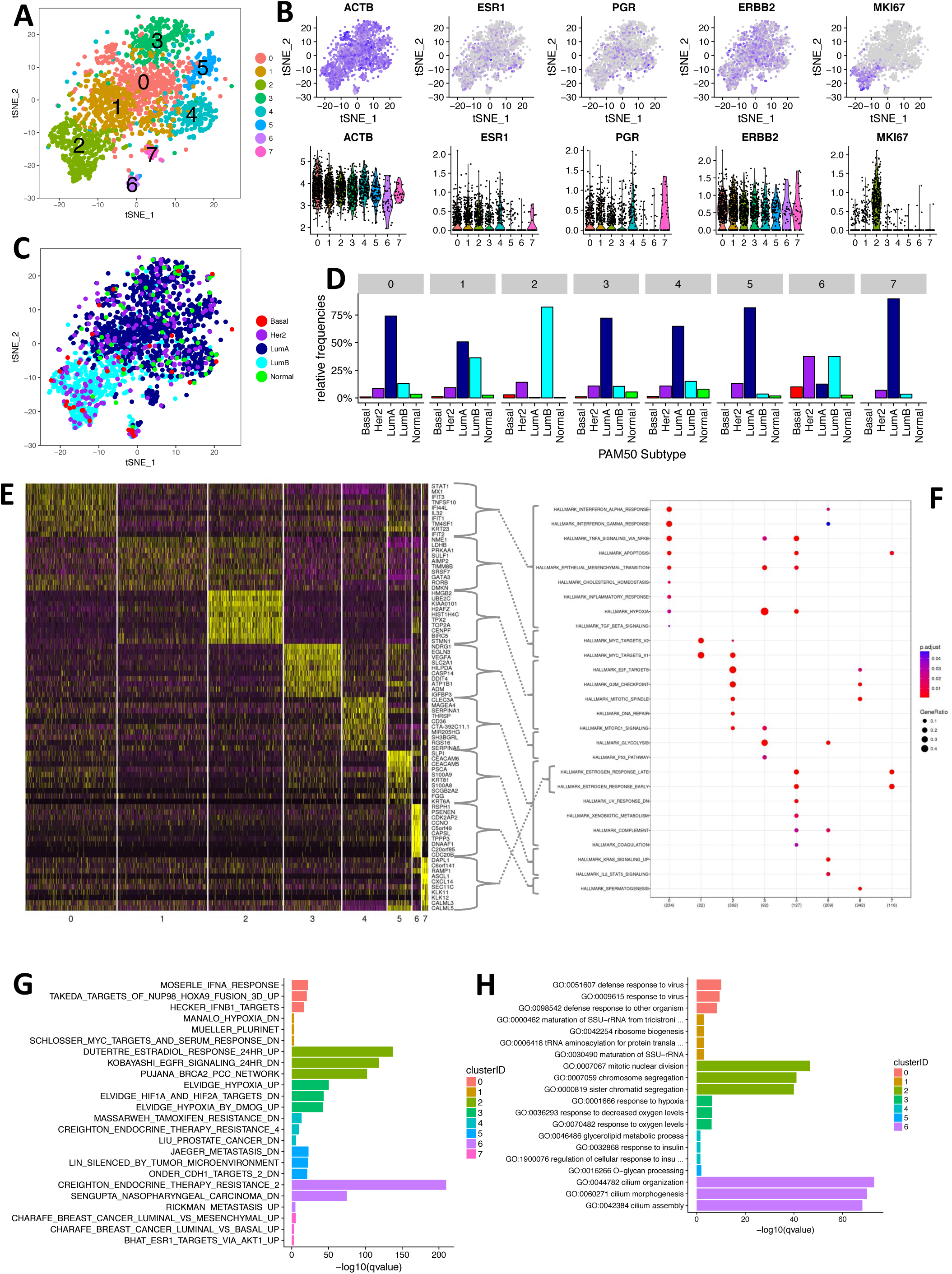
Unsupervised clustering of single-cell RNA-Seq identifies sub-populations of functional heterogeneity and putative drug resistant cells in clinical models of ER+ breast cancer. **A)** t-SNE visualization showing the cluster membership of each cell in PDX-HCI003. **B)** Expression level of selected genes in each cell superimposed on the t-SNE (top) and as violin plots grouped into the unsupervised cluster identities (bottom). **C)** t-SNE showing the PAM50 subtype classification for each cell. **D)** Relative frequency ofPAM50 subtypes in each of the eight cell clusters**. E)** Heatmap of the top-10 marker genes that are differentially expressed in each of the eight clusters. **F)** The MSigDB Hallmark gene-sets that are significantly (q-value<0.05) functionally enriched in each cluster. All cluster positive marker genes were used as input for each cluster. **G and H)** The top-3 MSigDB “C2” gene-sets **(G)** and Gene Ontology molecular function **(H)** annotations that are significantly (q-value<0.05) functionally enriched in each cluster. Cluster unique positive marker genes were used as input for each cluster.

To explore the transcriptional and functional properties of these cell populations we identified the genes that were positively enriched in each (figure 4E) and used gene enrichment analysis to assign function (figures 4F-4H). Figure 4F shows that we can identify cell clusters that have shared and unique functional signatures. For instance: interferon signaling is enriched in both clusters 0 and 5; E2F targets, G2M checkpoint and mitotic spindle associated genes are shared between clusters 2 and 6; and estrogen response is enriched in both 4 and 7. In contrast, functional properties such as: cholesterol homeostasis; DNA repair; P53 pathway; xenobiotic metabolism and KRAS signaling are restricted to specific sub-populations. This starts to highlight the complex functional heterogeneity present within the tumour cells even when using a broad set of hallmark functional classifications (Liberzon et al., 2015).

To refine our view of the functional heterogeneity a similar analysis was run across the MSigDB C2 gene-sets and the gene ontology biological process annotations. To focus on functional properties that are characteristic of each individual cell subpopulation, a set of marker genes that are unique to each cluster was used. A summary of the most significant results is shown in figures 4G and 4H. As expected, we find that a number of similar functions are consistent with those in 4F, such as interferon signaling, cell cycle and hypoxia enrichment in cell clusters 0, 2, and 3, respectively. However, this approach also allows the identification of properties central to the biology of ER+ breast cancers, such as: growth factor signalling; epithelial differentiation; estradiol response; metastasis and luminal tumours (figure 4G). In particular, cells are identified with gene-signatures related to both endocrine treatment sensitivity (e.g., cluster 4 identifies cells that highly express genes that are down-regulated upon acquired tamoxifen resistance [MASSARWEH_TAMOXIFEN_RESISTANCE_DN]) and resistance (e.g., cluster 6 shows a very clear enrichment for genes that are associated with *de-novo* tamoxifen resistance [CREIGHTON_ENDOCRINE_RESISTANCE_2]). The cells in cluster 6 also show a strong enrichment for genes that are involved in primary cilia organization and assembly, identifying a potential link between a small subpopulation of ciliated tumour cells and *de-novo* therapy resistance. Both these clusters of possible treatment resistance also show decreased expression of ESR1 and PGR (figure 4B) and increased expression of SLPI, a known driver of metastasis in breast cancer (Wagenblast et al., 2015).

### Single-cell regulatory analysis recapitulates sub-populations of putative drug resistant cells in a treatment naïve PDX of ER+ breast cancer

To further explore the intra-tumour heterogeneity of cells that are potentially sensitive and denovo resistant to endocrine therapies a set of 7 endocrine therapy derived gene signatures (Creighton et al., 2008; Massarweh et al., 2008) were applied to each cluster and cell. Clusters 2 and both show enrichment for genes that are down regulated upon resistance to Tamoxifen, indicating that these cells may be most sensitive to Tamoxifen treatment (figure 5A). In contrast, clusters 0 and 5 share enrichment for the same transcriptional signatures related to endocrine therapy resistance, with cluster 6 uniquely enriched for the “group 2” de-novo resistance genes.

**Figure 5.**
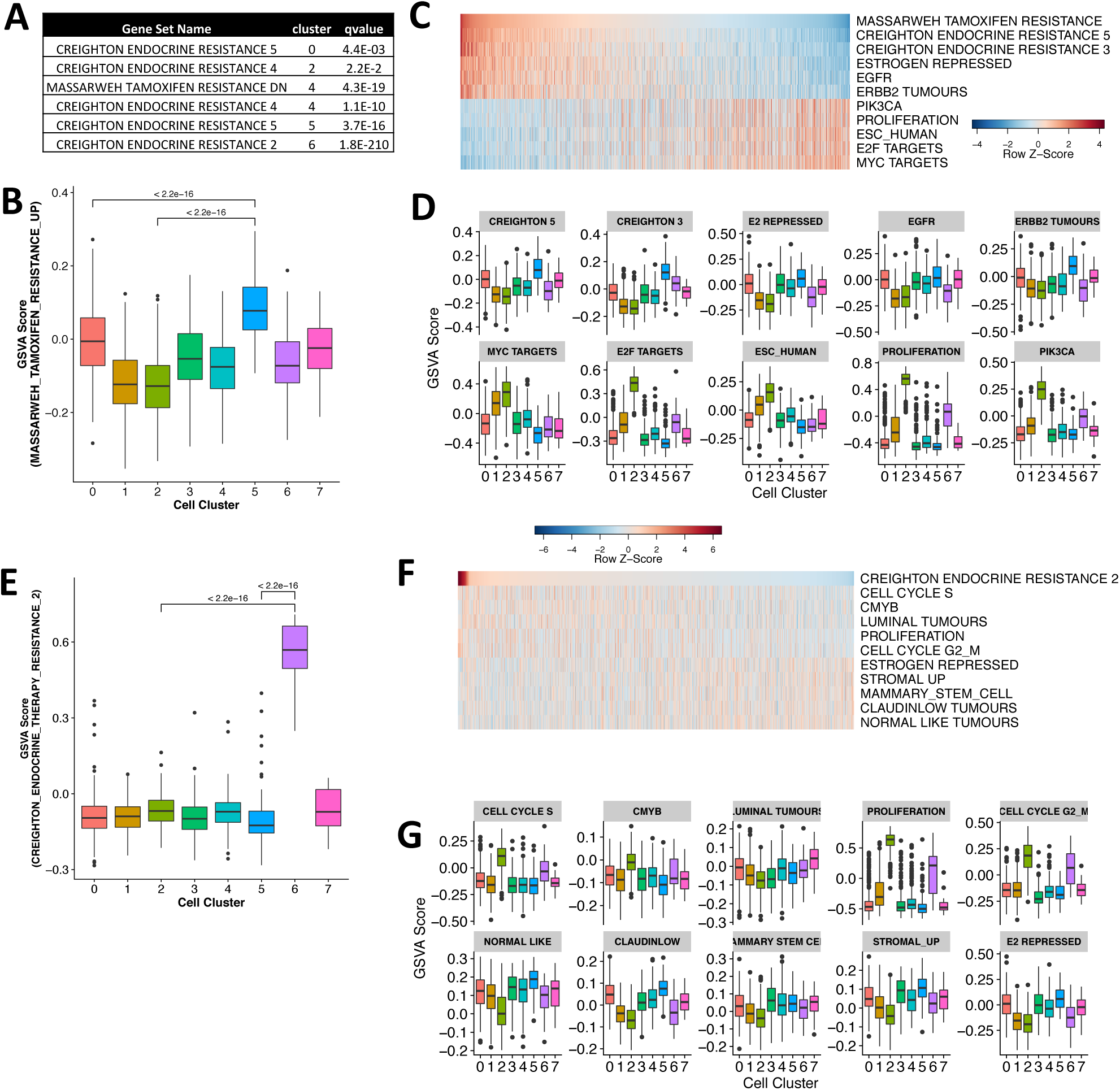
Single-cell regulatory analysis identifies distinct sub-populations and transcriptional drivers of putative drug resistant cells in a treatment naïve PDX of ER+ breast cancer. **A)** Per-cluster functional enrichment statistics for endocrine therapy treatment gene signatures. **B)** Boxplots showing the GSVA signature scores of all cells in each cell cluster from genes that are up-regulated in an MCF7 xenograft model of tamoxifen resistance. P-values calculated using Wilcox rank sum test. **C)** Heatmap of row-scaled GSVA scores from a non-redundant set of the top-5 positively (top) and negatively (bottom) correlated transcriptional signatures with the cellular enrichment for tamoxifen resistant gene-signature (Pearson correlation coefficient). **D)** Boxplots showing the GSVA scores for cells in each cluster for the correlated transcriptional signatures identified in C. **E)** Boxplots showing the GSVA signature scores of all cells in each cell cluster from genes that are up-regulated in an MCF7 xenograft model of *de-novo* tamoxifen and Gefitinib resistance. P-values calculated using Wilcox rank sum test. **F)** Heatmap of row-scaled GSVA scores from a non-redundant set of the top-5 positively (top) and negatively (bottom) correlated transcriptional signatures “Creighton therapy resistant 2” resistant gene-signature shown in E (Pearson correlation coefficient).

To complement this approach we generated GSVA functional signature scores for each individual cell rather than clusters, using a selected set of important breast cancer phenotypes (including the 7 endocrine therapy resistance signatures). This approach identified cells within cluster 5 as having a large significant enrichment (p<2.2e-16) for genes that are up-regulated in a xenograft model of tamoxifen resistance (Massarweh et al., 2008) (figure 5B). The gene signatures that most correlate with these were related to: two other endocrine treatment resistant models (CREIGHTON groups 3 and 5); estrogen repressed genes; EGFR signaling; and ERBB2 tumours (figures 5C and 5D). Interestingly, there was also a clear negative correlation with proliferative cells and those that show repression of Hallmark MYC transcriptional target genes (figure 5C and 5D). Further, we again identify cells within cluster 6 as a distinct group of cells with very clear enriched transcriptional properties of *de-novo* endocrine therapy resistance (figure 5E). Signature correlation across all cells highlighted moderate (<0.2) positive correlations for cell-cycle, proliferation, luminal tumours and the CMYB transcriptional program (figure 5F). In contrast, modest (>-0.16) negative correlations with mammary stem cell properties and the more mesenchymal normal-like and claudin-low tumour types (figure 5F) were found. These signature correlations were, however, much lower than those identified for the cluster 5 cells, where the largest positive and negative correlations were 0.9 and -0.58, respectively, compared to 0.2 and - 0.16 in the cluster 6 analysis. We next hypothesized that identification of the key transcriptional drivers of these cells may provide further insight into the mechanisms of transcriptional heterogeneity.

### Single-cell gene regulatory networks can identify transcriptional drivers of putative drug resistant cells

SCENIC (Aibar et al., 2017) was used to build gene regulatory networks and identify cis-regulatory motifs that are active in each of the individual tumour cells. This allowed us to identify active transcription factor “regulons” and generate a high-resolution transcriptional map of 181 transcriptional drivers of intra-tumour heterogeneity (figure 6A). We first looked to see if this method could identify the transcriptional drivers of proliferation present in the luminal B and cluster 2 cells (figure 6A). Indeed, E2F transcriptional activators (E2F1 and E2F2) and co-factors (TFDP1) were active in these cells, along with other components of cell cycle control (MYBL1, MYBL2, BRCA1). An E2F transcriptional repressor (E2F8) and factors related to stem cell maintenance (HMGB3, POU2F1) were also active in these cells.

**Figure 6.**
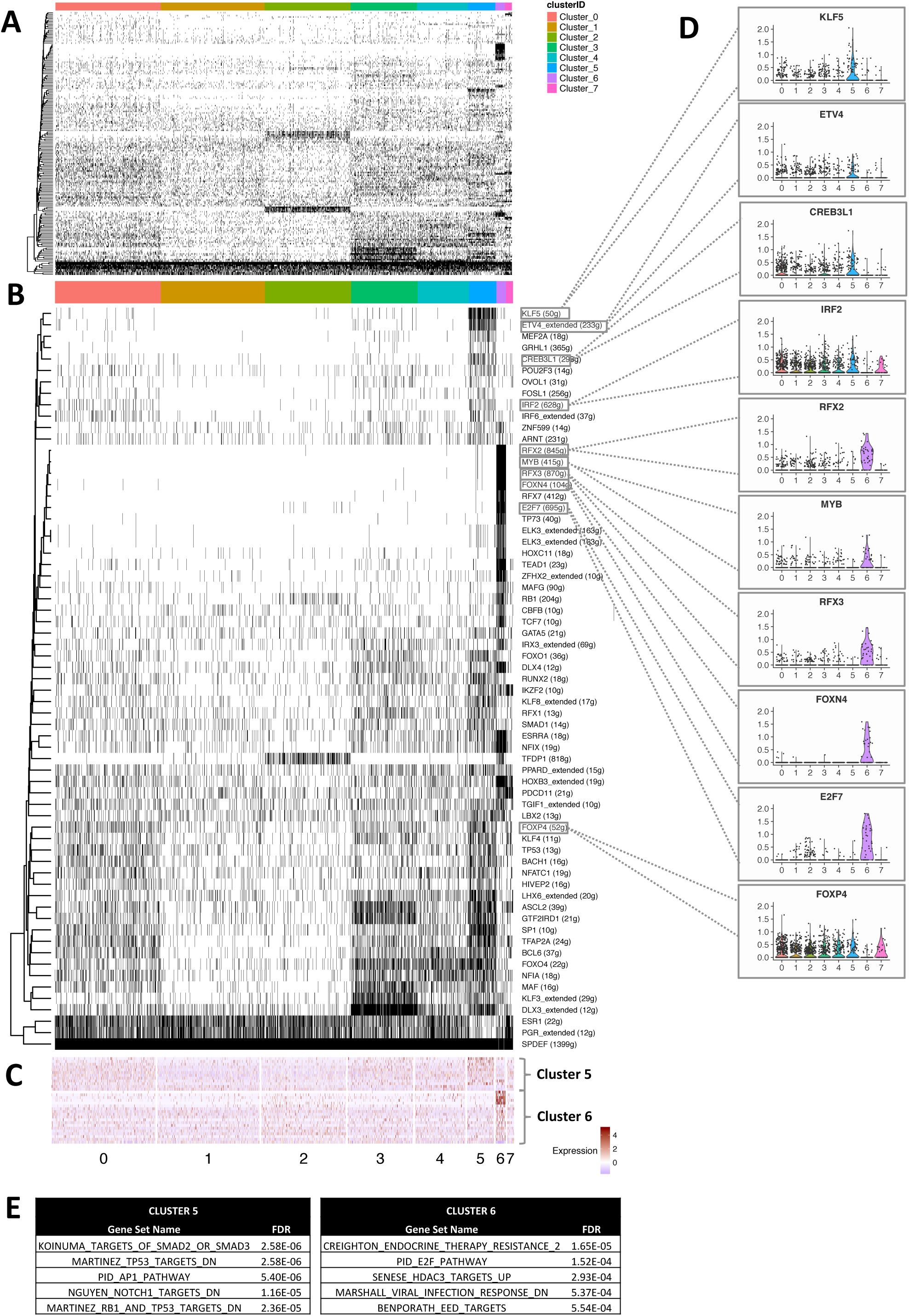
Single-cell gene regulatory network analysis can identify transcriptional drivers of putative treatment resistant breast cancer cells. **A)** Heatmap of active transcription factor regulons in each cell. **B)** Heatmap of transcription factor regulons with activity that is significantly enriched or depleted in cells from either cluster 5 or 6. (chi-square test p<0.001). **C)** Gene expression level of transcription factors, shown in B, that are also significantly enriched or depleted for expression in cell cluster 5 or 6. **D)** The single-cell gene expression levels of the transcription factors that drive the top-5 transcription factor regulons identified as significantly enriched for activity in clusters 5 or 6. **E)** The top-5 gene signatures that are enriched for the significantly activated transcription factors in cells within clusters 5 and 6.

We next identified the TFs that were significantly (chi-square p<0.001, after Bonferroni correction) activated or repressed in either of the two putative endocrine therapy resistant clusters (figure 6B). This resulted in 38 and 24 significantly activated TFs in cell clusters 5 and 6, respectively. Although the majority of differences were due to increased activation, we did also see significant reduction of both the ESR1 (Bonferroni p=1.95e-12) and PGR (Bonferroni p=4.88e-13) gene regulatory networks in the cells from cluster 5, mirroring the reduction in gene expression seen in figure 4B. With regards to the activated regulons, there was a clear, distinct group of factors that were almost exclusively active in the cells from cluster 6. This pattern was also reflected in the significantly increased level of expression, in cluster 6 when compared to all other cells, of a number of the transcripts that code for these factors (figures 6C and 6D). These included: E2F7, a transcriptional repressor that has been shown to have a role in endocrine resistance and predicting relapse in breast cancer (Browne et al., 2018; Chu et al., 2015); RFX3, a master regulator of cilia formation, which has a role in invasion (Légaré, Chabot, & Basik, 2017); as well as MYB (a cell cycle control factor) and FOXN4.

A similar analysis of the cells from cluster 5 identified significant activation and increased expression of KLF5 (figures 6B and 6D). These cells are also enriched for expression of transcription factors that have a role in the SMAD2/3 signalling pathway and the AP-1 transcriptional network (figure 6E). There was also a more complex overlapping pattern of transcriptional regulation with the hypoxic cells in cluster 3. For instance, ASCL2 is thought to maintain a role in maintaining selfrenewal in myogeneis (Wang et al., 2017) and has been shown to play a role in self-renewal of progenitor cells in colon cancer in conjunction with the YAP/KLF5 transcriptional axis (Wei et al.,2017). These results highlight how this approach can be used to uncover potentially novel transcriptional drivers of endocrine resistance and cancer cell self-renewal, and also highlight the complex heterogenous patterns of transcriptional regulation between drug resistant and hypoxic cells.

## Discussion

We describe the transcriptional heterogeneity present within the neoplastic cells from a treatment naïve PDX model of ER+ breast cancer as well as cancer cells from a diverse panel of breast cancer cell-lines. Using two complementary methods for transcriptionally profiling singlecells we have shown that breast cancer cells show significant transcriptional heterogeneity and that have heterogenous distributions of intrinsic PAM50 molecular subtype. Surprisingly, we were able to identify significant transcriptional and subtype heterogeneity even in clonal cell-lines that are commonly thought to be homogenous models of disease. We showed that in both the sensitive qPCR method and our high-throughput analyses using the 10X Chromium system that the transcriptional profiles of the individual cells for each subtype have similar transcriptional properties to those of bulk tumours (Curtis et al., 2012). For instance, the luminal B and basal-like cells show a similar increased proliferative nature to that seen in bulk tumours, whereas cells classified as luminal A or normal-like have little expression of proliferative marker genes, such as MKI67 (figures 2B-2E and 2H).

A potentially important consideration when assessing molecular subtype at cellular level instead of a bulk tumour is that we begin to see conflicting classifications with regards to the largest proportion of cells of a specific subtype and the overall tumour classification. For instance, the SKBR3 cell-line is classified as Her2-enriched at a bulk level, whereas in our single-cell analysis the Her2-enriched cells are the second largest group. Similarly, in our high-throughput dataset of the luminal B ER+ PDX model (DeRose et al., 2011), the luminal A cell proportion is largest. Although a much larger analysis of intra-tumour subtype heterogeneity will be required to gain a systematic comparison and assessment of bulk vs cellular subtype calls on clinical outcome, it observations indicate that it’ s likely to add an additional dimension of complexity to disease classification.

Hormone receptor signalling is a vital component of normal breast development and cancer. Our transcriptional profiling of ∼3500 cells from two models of ER+ breast cancer has provided, for the first time, an opportunity to explore the heterogeneity present in ER cis-regulatory signalling. We have shown that ER expression alone, or in combination with FOXA1, isn’t sufficient to maximally activate ER target genes within individual cells (figure 3). It is only when cells that co-express PR, a co-regulator and down-stream target of ER (Mohammed et al., 2015), are considered that we get the largest activation of direct ER targets (figure 3E). Interestingly, a similar level of activation of ER targets is also seen for cells lacking FOXA1 expression (i.e., ER+/FOXA1-/PR+). Although this is initially somewhat counter-intuitive, as FOXA1 has been shown to be necessary for canonical ER activation (Hurtado et al., 2011), it may be that the pioneer role of FOXA1 in ER transcription is a fleeting event and is not required to maintain the ER transcriptional process, with co-expression of co-regulatory factors more important.

Using unsupervised computational clustering approaches in the MCF7 cell-line and HCI003 PDX model, we observe clear transcriptional features of many of the hallmarks of cancer between clusters (Hanahan & Weinberg, 2011). These manifest in complex patterns of cellular subpopulations that have unique and common functional characteristics, such as: inflammation; proliferation; hypoxia, angiogenesis and glycolysis; growth-factor, TGF-β signalling and EMT; and invasion. This adds to previous studies showing that PDX models are valuable pre-clinical models of broadly conserved features of tumour heterogeneity (Bruna et al., 2016; DeRose et al., 2011).

Predicting the efficacy of treatment response and prognosis is essential in cancer treatment. Endocrine therapies have been revolutionary in the treatment of luminal breast cancers through their targeted control of the ER transcriptional response. They are, however, prone to resistance and relapse. One avenue for relapse is that cells within a tumour may have *de-novo* resistance to treatment and may ultimately be refractory to treatment. Importantly, we show that it is possible to identify cells, within the same tumour, that are potentially both sensitive and resistant to standard endocrine treatments (figure 4 and 5). Surprisingly, the proliferative cells appear to have transcriptional properties that are most similar to tumours that are sensitive to treatment, whereas the putative resistant cells in clusters 5 and 6 have very low and intermediate proliferative signatures, respectively. Future studies will assess the functional significance of these populations to endocrine response in PDX models.

Previous studies have identified a link between endocrine resistance and acquisition of a Her2-enriched transcriptional phenotype (Creighton et al., 2008; Massarweh et al., 2008). Similarly, we show that cells from the PDX model that are classified as the HER2+ subtype are also depleted for genes that are down-regulated in tamoxifen resistant tumours (Massarweh et al., 2008). However, association with a single subtype does not appear to be the only factor driving endocrine resistance as we observe that the cells in both of our de-novo resistant clusters contain a mix of subtypes. In particular, the cluster 6 sub-population has a striking mix of subtypes (figure 4D). This “Creighton 2” transcriptional signature was also uniquely enriched in both the basal-like and her2 cells from the PDX model. One intriguing explanation is that we are seeing the emergence of basal-like and hormone receptor negative cells that are also associated with *de-novo* treatment resistance. Previous work has reported the existence in luminal breast cancers of rare, stem-like, hormone receptor negative cells that express a basal cytokeratin (CK5+) (Kabos et al., 2011). Indeed, both sub-populations have significant reduction in ER and PR gene expression (figure 4B) and reduced activation of the ESR1 transcriptional regulons (figure 6B). In addition, cluster 5 has unique expression of both basal (KRT6A) and cancer stem cell (ALDH1A3) markers (Kabos et al., 2011) adding further weight to this possibility.

Invasion is a key hallmark of cancer that drives metastasis (Hanahan & Weinberg, 2011). An interesting feature of the cells in cluster 6 was the highly significant enrichment for genes involved in cilia organisation, morphogenesis and assembly. Recent work has highlighted a role for primary cilia in driving invasion in breast cancer, especially within the triple-negative subtype (Legare et al.,2017). In particular, they identify the role of SPEN and RFX3 as drivers of invasive cilia formation, both of which we identified as being uniquely and significantly expressed in the cluster 6 cells (figure 6D). A known marker of lung metastasis, SLPI (Wagenblast et al., 2015), is also identified as differentially expressed in both clusters 5 and 6 (figure 6E), drawing a further connection between treatment resistance and invasive potential. Using gene regulatory networks we identified potential transcriptional drivers of our cell populations associated with de-novo resistance (figures 6A and 6B). Significant enrichment was found for components of the SMAD/TGFB and AP1 transcriptional pathways (figure 6E), which have both been identified as playing a role in endocrine resistance (Browne et al., 2018; He et al., 2018). A number of candidate transcription factors were identified for the control of the putative resistant cells that express core components of primary cilia (figure 6B). These included RFX3, which is a master regulator of cilia formation (Legare et al., 2017), and also MYB and FOXN4, which have roles in ciliagenesis and promoting S-phase of the cell cycle (Campbell, Quigley, & Kintner, 2017; Tan et al., 2013). Further, the transcriptional repressor, E2F7, was significantly expressed and transcriptionally activated in the cilia associated cells. This E2F factor has an emerging role in endocrine resistance (Browne et al.,2018) and it’ s association with a role in primary cilia formation and cancer progression and relapse in luminal breast cancers deserves further investigation. The apparent rarity of these cells indicates that high-throughput single-cell methods would be ideally suited for studying these cells.

In conclusion, we have shown that intrinsic molecular subtypes can be assigned to individual breast cancer cells and a structure and organisation to cellular heterogeneity within breast cancer is emerging. We look forward applying these methods to larger sets of PDX models and clinical breast cancers. We have also identified putative transcriptional drivers of cellular sub-populations in a treatment naïve PDX tumour, that may have *de-novo* resistance to endocrine therapy and an unexpected association with primary cilia formation. These results highlight the power and application of high-resolution single-cell datasets to identify transcriptional heterogeneity of potential clinical relevance and ultimately track these cells during treatment.

## Materials and Methods

### Cell lines (Fluidigm qPCR)

MCF7 cell line was a gift from Michigan Cancer Foundation (Michigan, USA). The MDA-MB-231 cell line was a gift from EG & G Mason Research Institute (Massachusetts, USA). All other cell lines were obtained from American Type Culture Collection. Cell lines were maintained in RPMI 1640 media (Gibco) supplemented with 1-% FBS (Gibco), 20mM HEPES (Gibco) and 0.25% insulin (Novo Nordisk). Cells were seeded at day 0 and harvested at day 2 at ∼80% confluence. Cell suspensions were pipetted several times to ensure a single-cell suspension, which was confirmed using the Countess Automated Cell Counter from Invitrogen.

### Single-cell capture and gene expression analysis (Fluidigm qPCR)

Average cell size was determined using Countess Automated Cell Counter. Cell size for each cell line was within the range suitable for the 10-17um Fluidigm Integrated Fluidic Circuit C1 chip. A single cell suspension of ∼200,000 cells/ml was prepared in standard cell culture medium. Cell capture was conducted using the Fluidigm C1 Single-cell Auto Prep System. Cells were processed as follows; briefly, cells were lysed, RNA extracted and reverse transcribed. Genes of interest were pre-amplified using Taqman gene expression assays (Sup Table 1) and analysed for gene expression using a 96.96 gene expression chip and the Fluidigm Biomark HD Gene Expression System. A ‘Tube Control’ of 1000 cells of each cell suspension was prepared in parallel (as for bulk samples below) with the Fluidigm C1 Single-cell Auto Prep System. A ‘Mixed Tube Control’ was prepared by mixing equal volumes of each cell line’ s Tube Control. The Tube Controls and the Mixed Tube Control were analysed on each gene expression chip. This served as quality control to enable confirmation of Taqman assay performance during single-cell analysis, even if single cells did not express a particular gene.

### Bulk RNA preparation and analysis (Fluidigm qPCR)

Cells were seeded as above for single-cell analysis and harvested using the miRNeasy Mini Kit (QIAGEN). The Transcriptor High Fidelity cDNA Synthesis Kit (Roche) was used with 4ug of RNA. Samples were amplified using the Applied Biosystems 7900HT Fast Real-Time PCR System combined with Taqman Gene Expression Master Mix (Life Technologies) and Taqman Gene Expression Assays.

### MCF7 Species mix sample preparation for 10X single-cell capture

The MCF-7 cell line was obtained from the American Type Cell Culture Collection (ATCC) and grown in RPMI-1640 supplemented with 10% (v/v) fetal bovine serum (FBS; Gibco ®) and human insulin (100U/mL; Novo Nordisk). The mouse MEC cell line Comma-Dβ was a gift from Joseph Jeffery (University of Massachusetts, Amherst, MA, USA). Comma-Dβ cells were maintained in DMEM/F12 media (Gibco®), supplemented with 2% FBS (Thermo-Scientific), 10 mM HEPES (Gibco ®), 5 ml penicillin/streptomycin (Gibco ®), 0.25% insulin (Novo Nordisk) and 5 ng ml^−1^ murine epidermal growth factor (mEGF) (Sigma). All cell lines were grown at 37°C in a 5% CO_2_ incubator. Each cell line was characterized by short tandem repeat analysis (STR) profiling using the PowerPlexR 18D System (Promega) and tested for mycoplasma contamination (MycoAlert™ Mycoplasma Detection kit, Lonza).

### Patient derived xenograft - animal procedures, surgery and tumour dissociation

All animal procedures were carried out in accordance with relevant national and international guidelines and according to the animal protocol approved by the Garvan/St Vincent’s Animal Ethics Committee (Animal ethics number 15/10). The PDX HCI-003 transplants were carried out by surgical injection via direct visualization into the fourth mammary fat pads of pre-pubescent NOD-scid IL2rγnull (NSG) mice.

**The** PDX tumor was processed into single cell suspensions single cell capture. Tumor dissociation into single cell suspension was carried out using the MACS human Tissue Dissociation Kit (Miltenyi Biotec, Australia) following the manufacturer’ s instructions. The sample was then resuspended in RPMI 1640 and filtered sequentially through 70 μM and 40 μM cell strainers (BD Falcon) and the resulting single cell suspension was centrifuged at 300 x g for 7 min. Cells were then resuspended in 1X BD Pharm Lyse™ lysing solution (555899, BD Biosciences) for 3 minutes at room temperature (RT) to lyse erythrocytes.

### Flow cytometry and FACS isolation

Cell sorting and live cell enrichment experiments were performed at the Garvan Institute Flow Cytometry Facility. FACS experiments were performed on a FACS AriaII sorter using the BD FACSorter software.

### Single-cell capture and sequencing (10X)

The PDX HCI-003 tumor model was processed into single cell suspensions as described previously. Single epithelial and stromal cells were loaded on a Chromium Single Cell Instrument (10x Genomics, Pleasanton, CA) to generate single cell GEMs. As per manufacturer instructions, approximately ∼8,700 cells were loaded per channel for a target of ∼5,000 cells (10x Genomics). The MCF7 and Comma-Dβ cell-lines were mixed in cell suspension with a 50:50 ratio and loaded on a Chromium Single Cell Instrument (10x Genomics, Pleasanton, CA) to generate single cell GEMs. As per manufacturer instructions, approximately ∼5,300 cells were loaded per channel for a target of ∼3,000 cells (10x Genomics).

Single cell cDNA and RNA-Seq libraries were prepared using the Chromium Single Cell 3’ Library and Gel Bead Kit v2 (10x Genomics) and the products were quantified on the 2100 Bioanalyzer DNA High Sensitivity chip (Agilent). Both single cell libraries were sequenced using the Illumina NextSeq 500 system using the following parameters: pair-end sequencing with single indexing, 26 cycles for Read1, 8 cycles for I7 Index Read and 98 cycles for Read2.

### Single-cell gene expression analysis (Fluidigm qPCR)

Single-cell gene expression data was normalised by calculating the arithmetic mean of the house keeping genes (HKGs) for each chip, the arithmetic mean of all these values combined and then subtracting the latter from the former and combing this value with the raw Ct value for each reaction. Unsupervised clustering analysis of the pre- and post-normalised data confirmed the effectiveness of the method (not shown) and normalisation was therefore assumed to be successful in removing any batch effect caused by the separate chips.

Log2Ex represents the transcript level above background expression. To transform the qPCR Ct values into Log2Ex a limit of detection (LoD) value is required (equation below). To identify this, the Fano factor (variance [σ^2^] divided by mean [μ]) was calculated for each assay across a range of LoD values (20-40). The minimum LoD value at which the Fano factor for all assays was greater than or equal to one was selected (Livak et al., 2013).

~~~
Log2Ex = LoD − Ct if Ct ≤ LoD
Log2Ex = 0 if Ct > LoD
~~~

Any cell not expressing both HKG (GAPDH or HPRT) or expressing either below three standard deviations of the mean was excluded (Treutlein et al., 2014). This resulted in 359 single cells with normalised expression data and passing QC filters.

### PAM50 molecular subtype classification (qPCR)

The majority of this analysis was performed using a modified version of the PAM50 subtype tool (https://genome.unc.edu/pubsup/breastGEO/) with some additional custom analysis methods, The subtype comparison with the highest correlation coefficient is then assigned to the cell.

### Single-cell RNA-Seq analysis

The Cell Ranger Single Cell Software (10X Genomics) was used to process raw bcl files to perform sample demultiplexing, barcode processing and single cell 3’ gene counting (https://software.10xgenomics.com/single-cell/overview/welcome). Each dataset was aligned to a combined human (hg19) and mouse (mm10) reference and only cells that were identified as aligned to human in the “filtered” output of the cellranger count module were used in the analysis. Seurat v2.0 (Satija et al., 2015) was used to log-normalise (using a scale factor of 10000) and scale and centre the count matrices from the 10X single-cell datasets. Cells with less than 500 genes detected per cell and greater than 20% mitochondrial gene expression were removed. PCA analysis was performed on the variable genes and the Jackstraw method, with a p-value threshold of 0.01, was used to identify the significant principal components. For each dataset, these principal components were used as input to the unsupervised clustering (using a resolution of 0.8) and tSNE analysis. Differential marker gene analysis was performed between cell groupings using the Seurat “bimod” method, with a log fold-change threshold of 0.1 and p-value significance threshold of 0.01.

### PAM50 molecular subtype classification (10X)

The log normalised expression data (from Seurat, before scaling and centering) was used as input to the subtype specific centering method for PAM50 molecular subtype classification (Zhao et al., 2015). Each of the datasets were processed separately, resulting in a PAM50 subtype classification for each individual cell. The “ER_pos” pre-computed subtypes were used for both the MCF7 and PDX-HCI003 datasets.

### Gene Signatures

Gene-sets from a number of sources were used: MSigDB C2 (Liberzon et al., 2011); MSigDB Hallmark (Liberzon et al., 2015); Gene Ontology (Biological Process); breast cancer related (Gatza, Silva, Parker, Fan, & Perou, 2014); endocrine resistance therapy (Creighton et al., 2008; Massarweh et al., 2008); ER cis-regulatory signalling (Hurtado et al., 2011)).

### Functional enrichment

Functional gene enrichment between cell groupings was carried out using the ClusterProfiler method (Yu, Wang, Han, & He, 2012), with a qvalue threshold of 0.05 considered as significant.

### Gene signature score analysis

The GSVA method (Hänzelmann, Castelo, & Guinney, 2013) was used to generate gene-set enrichment scores for each individual cell using a manually curated set of gene-sets specifically related to breast cancer and ER+ disease biology.

### Transcription factor gene regulatory network analysis

The SCENIC method (Aibar et al., 2017) was used, with default filtering thresholds, to build gene regulatory networks and identify active transcription factor regulons within each cell.

